# Structural basis for functional interactions in dimers of SLC26 transporters

**DOI:** 10.1101/518209

**Authors:** Yung-Ning Chang, Eva A. Jaumann, Katrin Reichel, Julia Hartmann, Dominik Oliver, Gerhard Hummer, Benesh Joseph, Eric R. Geertsma

**Author notes:** these authors contributed equally (Yung-Ning Chang, Eva A. Jaumann, Katrin Reichel). to whom correspondence should be addressed: ERG; BJ; GH.

## Abstract

The SLC26 family of transporters maintains anion equilibria in all kingdoms of life. The family shares a 7 + 7 transmembrane segments inverted repeat architecture with the SLC4 and SLC23 families, but holds a regulatory STAS domain in addition. While the only experimental SLC26 structure is monomeric, SLC26 proteins form structural and functional dimers in the lipid membrane. Here we resolve the structure of an SLC26 dimer embedded in a lipid membrane and characterize its functional relevance by combining PELDOR distance measurements and biochemical studies with MD simulations and spin-label ensemble refinement. Our structural model reveals a unique interface different from the SLC4 and SLC23 families. The functionally relevant STAS domain exerts a stabilizing effect on regions central in this dimer. Characterization of heterodimers indicates that protomers in the dimer functionally interact. The combined structural and functional data define the framework for a mechanistic understanding of functional cooperativity in SLC26 dimers.

## Introduction

The solute carrier family 26 (SLC26), also known as the sulfate permease (SulP) family, facilitates the transport of a broad variety of organic and inorganic anions^1^. Members of this family are found in all kingdoms of life and operate predominantly as secondary transporters (symporters and exchangers)^2, 3, 4^. As an exception, prestin (SLC26A5) functions as a voltage-sensitive motor protein that evokes robust length changes in outer hair cells and thereby contributes to cochlear amplification^5, 6^. The relevance of the SLC26 family in maintaining anion equilibria is underlined by the causative role of mammalian SLC26 proteins in diseases such as congenital chloride diarrhea^7^ and cytotoxic brain edema^8^.

SLC26 proteins are composed of a membrane-inserted transport domain and a carboxyl-terminal cytoplasmic STAS (sulfate transporter and anti-sigma factor antagonist) domain. The SLC26-STAS domain is relevant for intracellular trafficking^9, 10^ and protein-protein interactions^11, 12, 13^. Its deletion impairs substrate transport by the membrane domain^4, 10, 14^. The crystal structure of SLC26Dg, a prokaryotic SLC26 protein from *Deinococcus geothermalis,* revealed a spatially separated membrane and STAS domain^4^. The SLC26Dg membrane domain holds two intertwined inverted repeats of seven transmembrane segments (TMs). Despite a poor sequence homology, the SLC26 family shares this 7-TM inverted repeat (7TMIR) architecture with the SLC4 and SLC23 families that transport bicarbonate and nucleobases plus vitamin C, respectively^15, 16, 17, 18, 19, 20, 21, 22^. The fourteen TMs are arranged in two subdomains: a compact core domain that holds the substrate binding site as inferred from the location of the nucleobases in the SLC23 crystal structures^20, 21, 22^, and an elongated gate domain that shields one side of the core domain. A mounting body of evidence^16, 19, 22, 23^ suggests that these proteins operate based on an elevator alternating-access mode of transport^24^ involving a rigid body translation-rotation of the core domain with respect to the gate domain.

Dimeric states have been previously observed for pro- and eukaryotic members of the SLC4^25, 27^, SLC23^28^, and SLC26^4, 29, 30, 31, 32^ families. Recent structures subsequently confirmed this oligomeric state for SLC4^17, 18, 19^ and SLC23^21, 22^ proteins and indicated that in both families the gate domains form the main interaction surface between protomers, though each family appears to hold a distinct dimer interface. As the crystal structure of SLC26Dg captured the protein in a monomeric state, the mode of interaction between SLC26 protomers has remained elusive. Interestingly, the protomers within the SLC26 dimer have been found to interact functionally^29, 33^ despite the presence of a complete translocation path in each individual protomer. Here we provide the first structural and mechanistic insights in the allosteric interactions between SLC26 protomers. We integrated pulsed electron-electron double resonance (PELDOR, also known as double electron-electron resonance or DEER) distance measurements and *in vitro* transport studies with structural modeling and refinement using MD simulation to determine the architecture of the membrane-embedded SLC26 dimer and characterize its functional relevance.

## Results

### Resolving the SLC26Dg dimer interface in the lipid membrane

To define the SLC26Dg dimer interface, we used interspin distance constraints derived from PELDOR experiments. Since SLC26Dg is monomeric in its detergent-solubilized state, we reconstituted spin-labeled protomers in lipid membranes to assure dimer formation^4^. As the gate domain in the SLC4^17, 18, 19^ and SLC23^21, 22^ families establishes the main protomer-protomer contacts in the membrane (Fig. 1a,b), we engineered spin-labels at 13 different positions of the termini of gate domain helices in SLC26Dg. By site-directed modification of single-cysteine mutants, we added the probe 1-oxyl-2,2,5,5-tetramethylpyrroline-3-methyl-methanethiosulfonate (MTSSL)^34^ (Fig. 1c). Size-exclusion chromatography and transport assays in proteoliposomes demonstrated that all mutants were folded well and active (Suppl. fig 1).

**Fig. 1.**
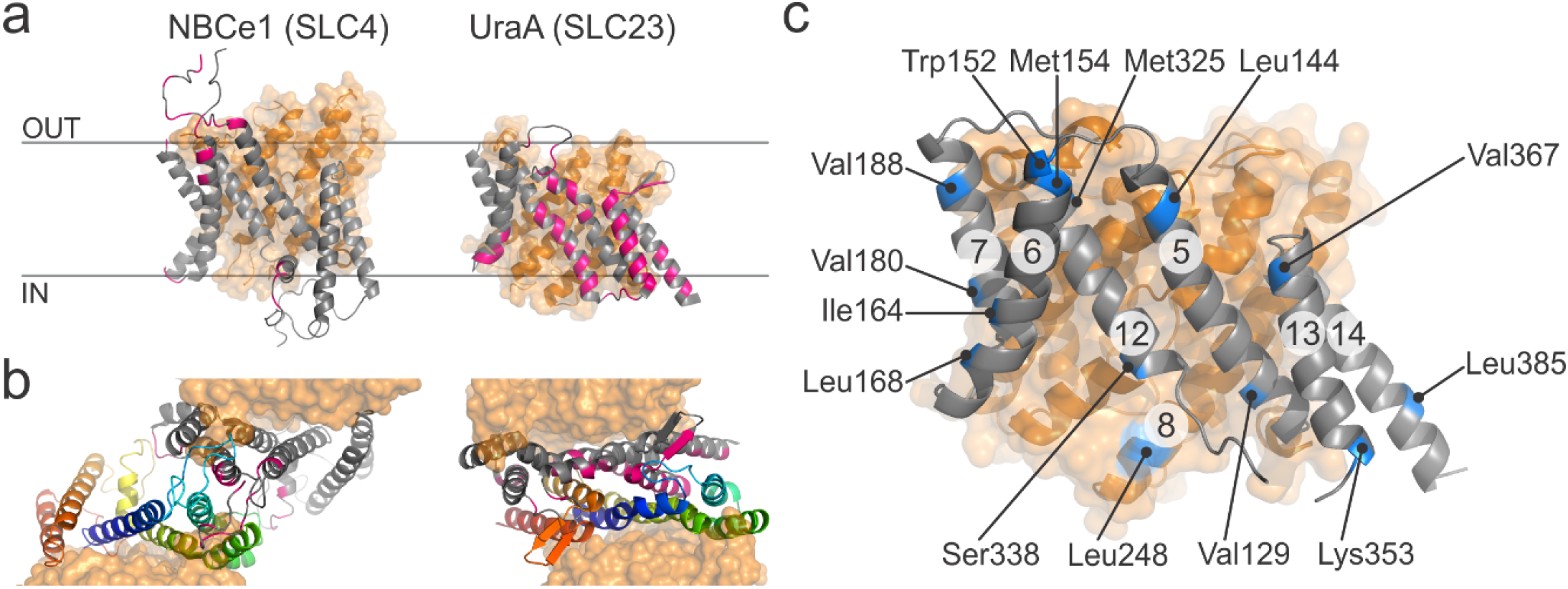
Dimer interfaces in 7TMIR proteins. **(a)** Side view of the membrane domains of NBCe1 (PDB: 6CAA) and UraA (PDB: 5XLS). Core and gate domain are colored orange and grey, respectively, with residues within 4 Å of the opposing protomer in pink. **(b)** Top views of the dimeric arrangements of NBCe1 and UraA. The gate domains of the two protomers follow a rainbow coloring scheme (blue-to-red for N-to-C direction). **(c)** Side view of the membrane domain of SLC26Dg (PDB: 5DA0). Residues mutated to cysteine for site-directed spin labeling are colored blue. The circled numbers indicate the respective TMs.

Systematic analysis of all positions led to the identification of three labeled positions, K353R1, V367R1, and L385R1 that gave well-defined interspin distance distributions centering around 4.4 ± 0.2, 3.9 ± 0.3, and 1.8 ± 0.1 nm, respectively (Fig. 2b-d). These positions are located in TM13 and TM14 and place this region in close proximity to the center of the SLC26Dg dimer interface. This particular dimer arrangement combined with the short phase memory time (*T_M_*) of the spins in membranes did not allow to accurately determine the long interspin distances between other helices (Suppl. fig. 2) An exponential decay was observed for the PELDOR measurement of the detergent-solubilized protein (Fig. 2a), supporting the notion that the identified region is part of the native SLC26Dg dimer interface formed in the lipid membrane. Given the spin-labeling efficiencies of 70-90%, the obtained modulation depths of the PELDOR time traces (12-24%), are in the range expected for a dimer, suggesting that the majority of the protomers in the membrane is part of a dimer.

**Fig. 2.**
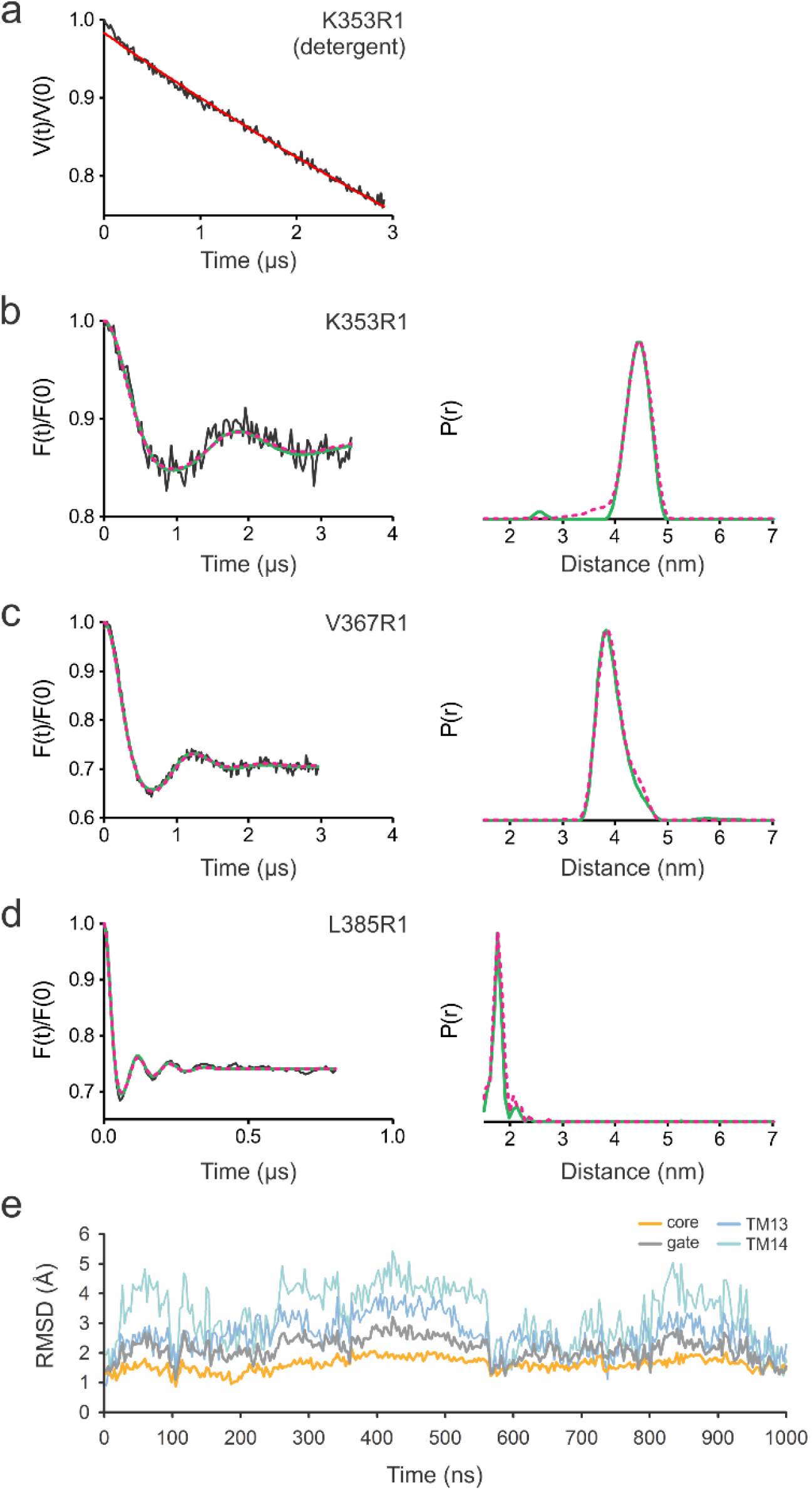
Interspin distances in the SLC26Dg dimer. **(a)** Primary PELDOR data of detergent-solubilized K353R1. **(b-d)** Left panels: Background corrected PELDOR time traces for membrane-reconstituted K353R1, V367R1, and L385R1 (black traces), overlaid with the fit from Tikhonov regularization (green), and forward-calculated PELDOR time traces from BioEn spin-label rotamer refinement of the MD simulation model (magenta, dashed; *θ* = 10). Right panels: distance distributions obtained by Tikhonov regularization (green), overlaid with the distance distributions resulting from BioEn analysis of the MD simulation model (magenta, dashed). Original PELDOR data in Suppl. Fig. 2. **(e)** C_α_-atom root-mean-squared-distance (RMSD) values of the core, gate, TM13 and 14 relative to the monomer crystal structure as a function of MD time (1 μs).

### Structural model of the SLC26Dg membrane dimer interface

On the basis of the PELDOR data and the SLC26Dg crystal structure, we constructed a dimer model. First, to obtain an equilibrated structure for rigid body docking, monomeric SLC26Dg without its carboxyl-terminal STAS domain was embedded in a POPC bilayer and submitted to 1 μs of MD simulations. We observed considerable flexibility of the gate domain in comparison to the core domain (Fig. 2e). In particular TM13 and TM14 exhibited significant motions in the monomer, in line with their suspected involvement in the membrane dimer interface. Due to the observed flexibility, we used a relaxed conformation, after 440 ns of MD, for docking. In a rigid-body search restricted by C2 symmetry with an axis normal to the membrane, we identified a candidate dimer structure for which the forward-calculated PELDOR traces matched the background-corrected time-domain data (Fig. 2b-d, left panels). An alternative rigid-body docking approach guided by the inferred distance distributions resulted in a very similar dimer model (Suppl. fig 3). The initial C2-symmetric MD dimer was then relaxed by additional MD simulation. The forward-calculated PELDOR traces for the relaxed dimer, after gentle spin-label rotamer refinement^35^, are in excellent agreement with the experimental data (Fig. 2b-d, left panels, Suppl. fig 4). The structure of this SLC26Dg membrane domain dimer model is shown in Fig. 3.

**Fig. 3.**
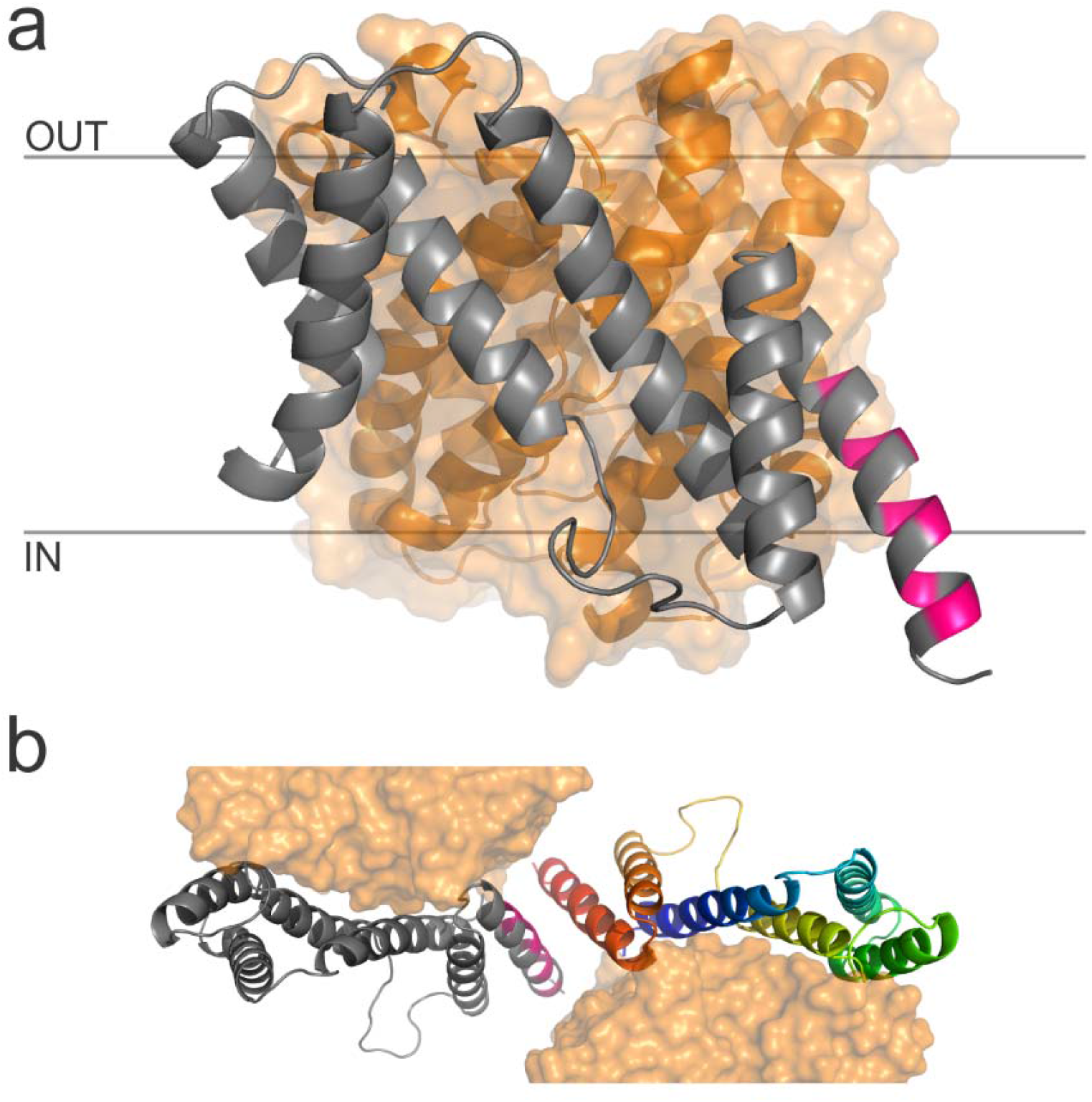
Model of the SLC26Dg dimer interface, **(a)** Side view of the SLC26Dg membrane domain in the same orientation as Fig. 1a. Core and gate domain are colored orange and grey, respectively, with residues within 4 Å of the opposing protomer in pink. **(b)** Top views of the dimeric arrangement of SLC26Dg. The gate domain of one of the protomers follows a rainbow coloring scheme (blue-to-red for N-to-C direction).

The model of the SLC26Dg dimer displays a protomer-protomer membrane interface that is remarkably different from the membrane interfaces observed for the SLC4 and SLC23 families, both in its location and in its size^17, 18, 19, 21, 22^. Whereas the membrane dimer interfaces of SLC4 and SLC23 proteins center around TM6, and TM5 plus TM12, respectively, the midpoint of the SLC26Dg dimer is TM14 on the opposite side of the gate domain. Furthermore, while the membrane dimer interface of SLC4 and SLC23 proteins involves extensive interactions covering approximately half and nearly the complete exposed membrane surface of their gate domains, respectively, the membrane interface of SLC26Dg is relatively small. Also in comparison to other oligomeric membrane proteins, the surface buried by dimerization of the membrane domain is modest^36^. This observation is in agreement with the complete absence of dimerization in detergent and suggests that other factors, such as subunit-bridging lipids or the cytoplasmic STAS domain may contribute to the stabilization of the dimeric state.

### STAS domain stabilizes regions central in the dimer

The cytoplasmic STAS domain is one of the major structural constituents that distinguishes the SLC26 family from the SLC4 and SLC23 families that do not hold carboxy-terminal domains^16^. While deletion of the STAS domain compromises the transport capacity of the SLC26Dg membrane domain, the structure of the membrane domain is not altered^4^. As the STAS domain immediately follows the central TM14, we further determined to what extent the STAS domain contributes to the dimer interface.

As evidenced from the PELDOR time trace for L385R1 in SLC26Dg^ΔSTAS^, deletion of the STAS domain did not affect the ability of the membrane domain to form dimers (Suppl. fig 5). STAS domain deletion resulted in a small increase in the mean L385R1 distance from 1.8 ± 0.1 to 2.1 ± 0.1 nm, indicating a modest rearrangement at the dimer center. In contrast, the analysis of SLC26Dg^ΔSTAS^-K353R1 and -V367R1 in TM13 revealed an enhanced flexibility and increased interspin distance (Suppl. fig 5). These observations indicate that the STAS domain stabilizes and affects regions in the center of the dimer.

### SLC26Dg dimer interface is representative for the SLC26 family

To further validate the SLC26Dg membrane dimer model and determine to what extent it represents the SLC26 family in general, we used oxidative cross-linking in biological membranes. Due to its central position we focused on TM14. Oxidative cross-linking of single-cysteine variants at several positions in TM14 of SLC26Dg lead to the appearance of a band with lower electrophoretic mobility (Fig. 4). We assign this band to SLC26Dg homodimers because an identical anomalous shift was observed on cross-linking in proteoliposomes (Suppl. fig 6). Crosslinks were observed for residues located at both ends of TM14, but not for residues facing the interior of the bilayer in line with a general lower reactivity of cysteines at this position^37, 38, 39^. The ability of cysteine residues in TM14 of SLC26Dg to form a disulfide bond with the opposing protomer further validates our SLC26Dg dimer model.

**Fig. 4.**
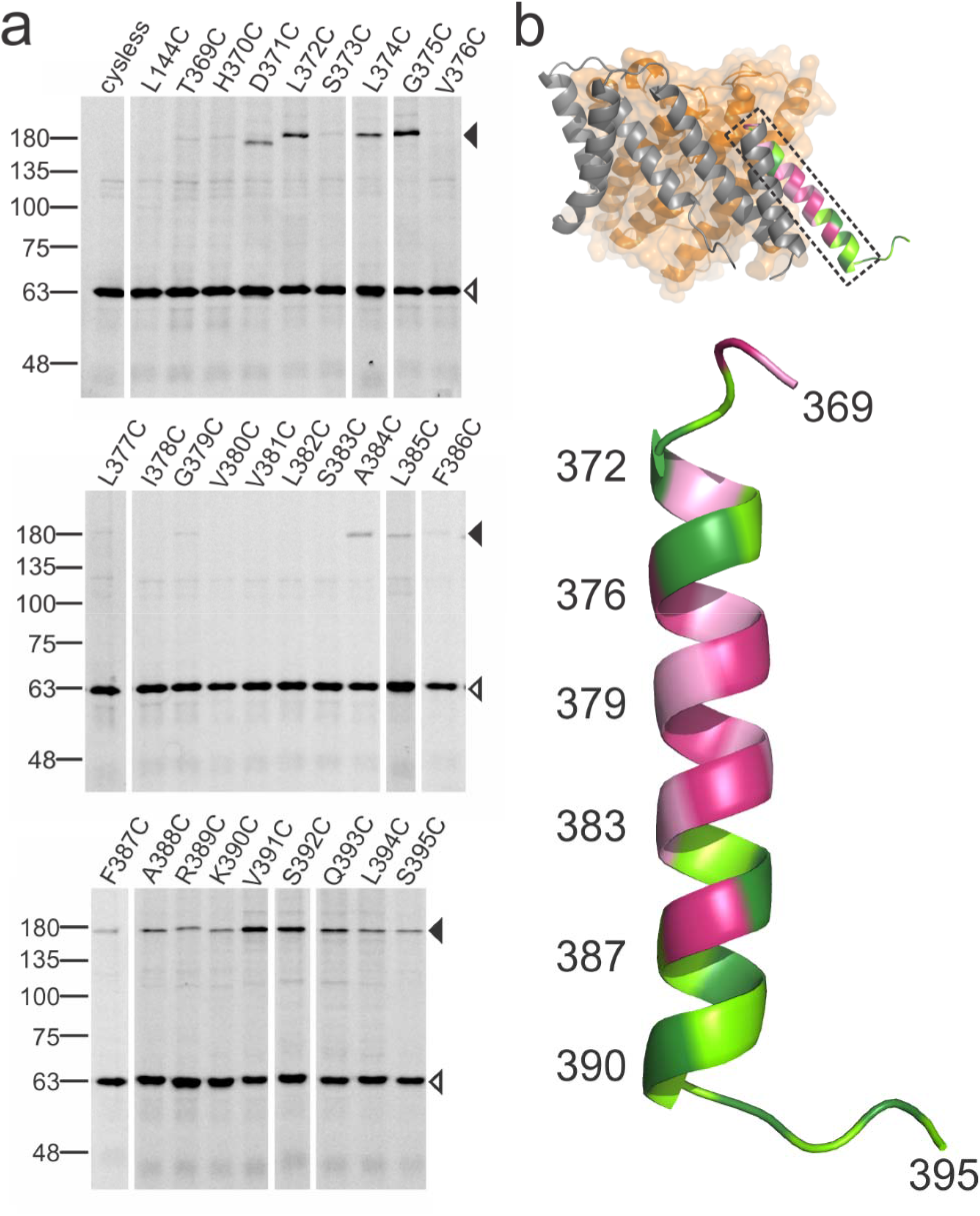
Oxidative cysteine crosslinking between TM14 of SLC26Dg. **(a)** In gel GFP fluorescence analysis of disrupted *E. coli* cells expressing single cysteine variants of SLC26Dg fused to superfolder-GFP. Following oxidative crosslinking, samples were analyzed by non-reducing SDS-PAGE. Cysteine-free SLC26Dg (cysless) and L144C (TM5) represent negative controls. Black and white arrows indicate dimeric and monomeric SLC26Dg. **(b)** Top panel: spatial orientation of TM14 in SLC26Dg. Lower panel: Enlargement of TM14 with positions susceptible to crosslinking colored in green, and non-susceptible positions in pink. Residues are alternatingly in light or dark color.

While the known dimer interfaces between SLC4 and SLC23 families differ greatly, a high degree of similarity is observed between members of the same family^17, 18, 19, 21, 22^. This suggests that the dimer interfaces for this fold are specific to a family. To test this, we used the same TM14 crosslinking approach on SLC26 proteins from *Sulfitobacter indolifex* and *Rattus norvegicus* (Suppl. fig 7–8). For both proteins, we observed the formation of TM14 disulfide cross-links between protomers, which provides first evidence that the membrane dimer interface may be very similar, if not conserved, throughout the SLC26 family.

### Functional relevance of the SLC26Dg dimer

The observation of a structural SLC26Dg dimer led us to ask whether this oligomeric state is important for function. As both protomers have independent binding sites and non-overlapping translocation paths, the relevance of the dimeric state is not evident. Functional interactions between protomers in oligomeric proteins can be revealed by mixing protomers with different functional characteristics and analyzing the resulting hetero-oligomers. We opted to create an inactive variant by locking the protein in the inward-facing conformation using disulfide crosslinking. Based on the crystal structure, we selected the extracellular side of TM1 (core) and TM5 (gate) as most suited concerning cross-linking efficiency and ability to lock the protein and prevent transport (Fig. 5a). Oxidative crosslinking of SLC26Dg-CL-I45C/A142C, hereafter named SLC26Dg-IL (inward-locked), resulted in a nearly complete shift in the electrophoretic mobility of the protein that could be restored by the addition of the reductant DTT (Fig. 5b). Likewise, fumarate transport of the cross-linked mutant in proteoliposomes was close to background activity, but could be fully recovered to wildtype activity by the addition of DTT, indicating that the protein was well-folded and reconstituted (Fig. 5c).

**Fig. 5.**
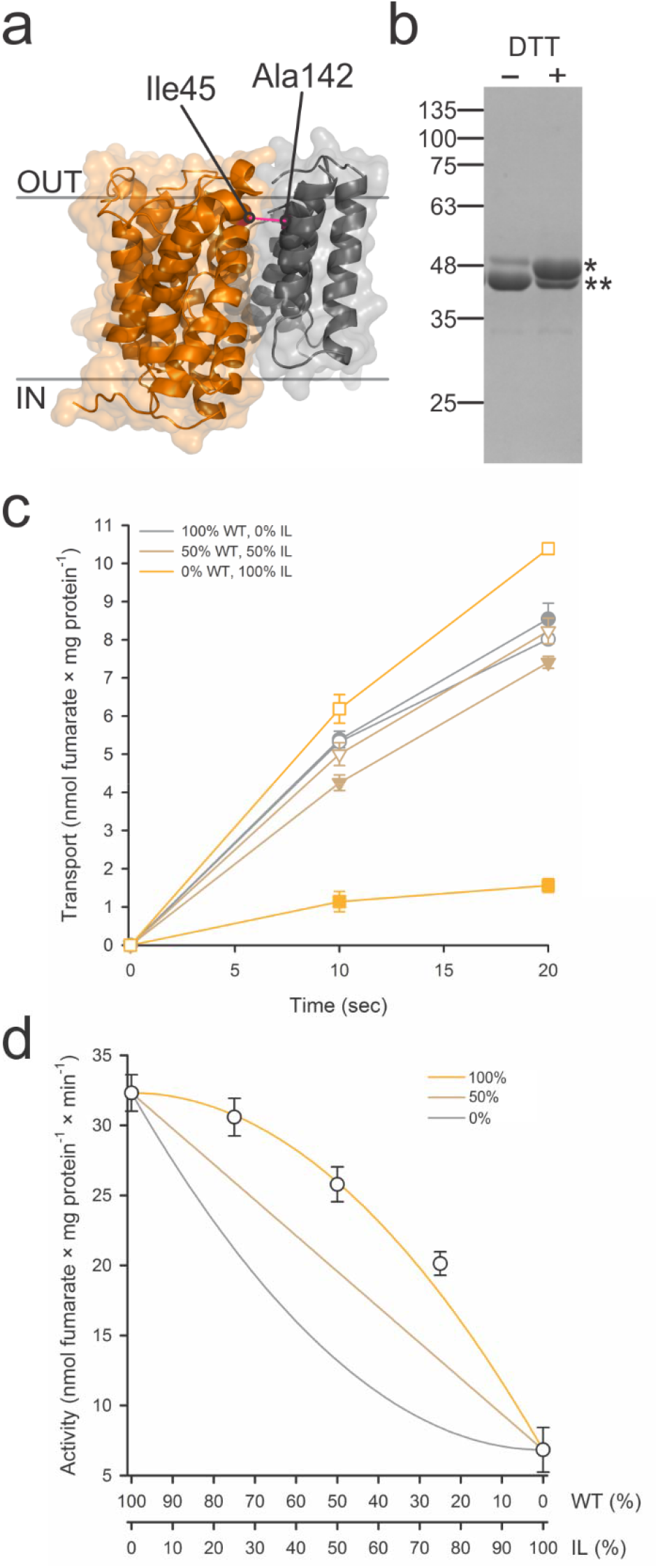
Generation and functional characterization of SLC26Dg-IL. **(a)** View in the plane of the membrane of the relative orientation of the cysteine mutants in the core (orange) and gate (grey) domain of SLC26Dg. **(b)** SDS-PAGE analysis of purified and crosslinked SLC26Dg-IL monomers in the absence and presence of DTT. Single and double stars indicate crosslinked and not-crosslinked protein. **(c)** Functional characterization of membrane-reconstituted and crosslinked SLC26Dg-IL (orange), wildtype SLC26Dg (grey), and both proteins mixed in equal ratio’s (brown). Closed and open symbols indicate the absence and presence of a pre-incubation step with DTT. **(d)** Initial transport rates of membrane-reconstituted and crosslinked samples containing wildtype and SLC26Dg-IL mixed in different ratio’s. Grey, brown, and orange curves indicate the anticipated curves assuming an activity of the heterodimers corresponding to 0, 50, and 100% of the wildtype homodimers. Data points represent mean and standard errors of three replicates.

As SLC26Dg is monomeric in detergent and dimerizes only after reconstitution in the lipid membrane, we achieved stochastic formation of heterodimers as demonstrated by the decreased TM14 cross-linking upon the addition of SLC26Dg-IL in a control experiment (Suppl. fig 9). Interestingly, the initial transport rates of proteoliposomes holding different ratios of wildtype and SLC26Dg-IL followed a positive quadratic relationship (Fig. 5d). The activity of the heterodimers exceeded the expected values for independent functioning of protomers, which is half the sum of the activities of the wildtype and SLC26Dg-IL homodimers (Fig. 5d, straight line). In fact, in the most parsimonious model for the quadratic dependence of the activity on the mixing ratio, only the SLC26Dg-IL homodimer is inactive and all other dimers have the same activity. This could imply that either only one protomer is active in a dimer or the activity of a wildtype protomer is doubled when paired with an inward-locked protomer. In any case, the strong coupling evident in this transport activity data is a strong indication that dimerization is functionally relevant.

## Discussion

The structural model of the SLC26Dg membrane domain dimer interface, based on EPR measurements on membrane-reconstituted protein and validated by cysteine cross-linking in biological membranes, places TM14 of the gate domain at the center of the SLC26Dg dimer. Further cross-linking studies on additional prokaryotic and mammalian homologues suggest that this interface is evolutionary conserved in the SLC26 family. The central position of the gate domain in the SLC26 membrane dimer interface corresponds well with the SLC4 and SLC23 families in which the gate domains also form the major contacts between the membrane domains^17, 18, 19, 21, 22^. However, although the dimer-interfaces seem conserved within a family, the regions involved in the protomer-protomer contacts seem to differ greatly amongst the three families.

It appears likely that these different dimeric arrangements represent stable constellations between which the protomers do not alternate during transport. The structures of dimeric SLC4 and SLC23 proteins in different conformations^17, 18, 19, 21, 22^ all hold identical, yet family-specific, contact surfaces. This is further supported by repeat-swap homology modeling of AE1, which indicated that no changes in the dimerization interface were required during the transition from the outward-facing structure to the inward-facing model^23^. Transitions between interfaces seem further unlikely due to the requirement for significant rearrangements in structural elements, such as the cytoplasmic region following TM12 in SLC4A1 and SLC4A4, which also appears stable and blocking alternative interfaces in SLC26Dg (Suppl. fig 10). Finally, our PELDOR data on SLC26Dg, especially the well-defined distance distributions for the interface region, strongly disagrees with a dynamic interface. A stable oligomer interface is in line with other observations on unrelated elevator proteins^40, 41^.

The buried surface resulting from dimerization in the SLC26Dg membrane domain amounts to ~350 Å^2^, which is small, not only in comparison to the SLC4 and SLC23 family whose membrane interfaces measure ~1000 Å^2^ and ~2000 Å^2^,^42^, respectively, but also in relation to other oligomeric membrane proteins^36^. It is likely that additional extrinsic factors contribute to extend and stabilize the SLC26 membrane dimer interface, e.g., interfacial lipids that were recently reported to stabilize an SLC23 dimer and other oligomeric membrane proteins^36, 43^. In this respect, the STAS domain appears to be relevant as well. The short linker region connecting TM14 and the STAS domain implies its close proximity to the membrane dimer interface. In addition, the STAS domain stabilizes TM13 and affects TM14 at the center of the dimer interface (Suppl. fig. 5). Given that isolated SLC26-STAS domains do not appear to form dimers^13, 44, 45^, we expect the STAS-mediated stabilization of the gate domain to result from a direct interaction between the STAS domain and the membrane domain. This interaction may also form the basis for the enhanced transport rates observed in the presence of the STAS domain^4, 10, 14^.

All 7TMIR proteins form structural dimers in the membrane, but the general relevance of this oligomeric state for their function is not clear. The available structures of the SLC4, SLC23, and SLC26 proteins all indicate that the complete substrate translocation path is contained within one protomer. This is supported by the recessive inheritance mode of SLC26-linked diseases^29, 46^ and further confirmed by the functional characterization of heterodimers composed of a wildtype and an inactive mutant protomer. Most of these heterodimers were found to be active for NBCe1^47^ (SLC4), UraA^22^ (SLC23) and SLC26Dg (SLC26, this work), though for UapA (SLC23) inactive heterodimers were observed as well^21^. With the exception of SLC26Dg, these studies were carried out in the context of whole cells, employed different mutations that inferred in diverse ways with substrate transport, and, in case of NBCe1 and UraA, involved the use of concatemeric constructs. This diversity in experimental approaches makes it difficult to precisely compare these data. Nevertheless, although the inferred specific activity of the heterodimers of NBCe1 (50% active) suggests that the protomers can operate independently, the apparent negative and positive dominance observed for UapA, and UraA plus SLC26Dg, respectively, indicates that the dimeric state may have a functional role as well. This notion is further supported by studies on rat prestin (SLC26A5) heterodimers composed of protomers that in the context of a homodimer hold a very different voltage-dependence of their non-linear capacitance^29^. For these heterodimers an intermediate phenotype was observed, suggesting a strong cooperative interaction in which the two protomers jointly determine the voltage-dependence of the conformational changes.

Though the mechanistic basis for functional interactions in 7TMIR dimers is currently unclear, important insights were obtained from the characterization of monomeric 7TMIR proteins. Monomeric variants of UraA, generated by the introduction of bulky residues at the dimer interface, bind substrate with wildtype affinities and are thus expected to be well-folded, but are incapable of facilitating transport^22^. In our study, we characterized the transport properties of individual protomers as well, but in the context of a dimer. These SLC26Dg-WT protomers, embedded in WT-IL heterodimers, are fully active. In fact, the WT-IL heterodimers have the same activity as wildtype homodimers. Together these observations highlight the relevance of the interaction between opposing gate domains for facilitating transport. This interaction may stabilize an essential conformation of the gate domain required for transport, as suggested previously^22^ and in line with our observation that the transport-incompetent SLC26Dg^ΔSTAS^ undergoes small rearrangements in the gate domain. Alternatively, the gate-gate domain interaction may provide a stable membrane-embedded scaffold that enables the vertical translation of the core domain and its anticipated deformation of the bilayer. In this context, the inward-locked SLC26Dg protomer may serve as an extended scaffold that fixates the gate domain even better in the membrane, providing a rational for the apparent increase in transport rate observed for the wildtype protomer in the heterodimer. Additional structures of dimeric 7TMIR proteins in multiple states will be required to further pinpoint the role of the gate domain.

Understanding the transport mechanism of 7TMIR proteins requires that proteins are not studied only as individual protomers, but also in the context of the dimer, their functionally relevant oligomeric state. Structures of SLC4 and SLC23 proteins have provided exceptional insight into protomer interactions by providing snapshots of dimeric constellations, but the structure of a dimeric SLC26 protein has been elusive. Here, we have determined the architecture of dimeric, membrane-embedded SLC26Dg using an integrated structural biology approach. The SLC26 dimer interface is unique and distinguishes itself from SLC4 and SLC23 proteins. We have demonstrated that the interface is not dynamic, though deletion of the carboxyl-terminal STAS domain, which impairs transport, introduces flexibility in an adjacent region. Finally, our heterodimer studies have underlined the functional significance of the dimer. Together these structural, dynamic, and functional characterizations provide the framework for further studies on the SLC26 family and offer novel mechanistic insights that may extend to other elevator proteins as well.

## Materials and Methods

### Site specific mutagenesis of SLC26 transporters

Cysteine residues were introduced into pINITcat-SLC26Dg by Quikchange mutagenesis or a two-step PCR method (mega-primer approach). Sequence-validated pINITcat-SLC26Dg variants were subsequently subcloned into pBXC3GH by FX cloning^48^ for protein expression and purification.

### Protein expression and purification

*E. coli* MC1061 containing pBXC3GH-SLC26Dg or the variants was cultivated in 9 L TB/ampicillin in a fermenter (Bioengineering). Cells were grown at 37 °C until an OD_600_ ≈ 2 was reached, after which the temperature was gradually decreased to 25 °C over the course of 1 h. Expression was induced by the addition of 0.005% (w/v) L-arabinose and continued for 16 h. Cell pellets were resuspended in 50 mM potassium phosphate (KPi), pH 7.5, 150 mM NaCl, and 1 mM MgSO4 and incubated for 1 h at 4 °C in the presence of 1 mg/mL lysozyme and traces of DNase I before disruption with an APV Gaulin/Manton homogenizer. The lysate was cleared by low-spin centrifugation, and membrane vesicles were obtained by ultracentrifugation. Vesicles were resuspended to 0.5 g/mL in 50 mM KPi, pH 7.5, 150 mM NaCl and 10% glycerol (buffer A). All subsequent steps were carried out at 4 °C. Membrane proteins were extracted for 1 h at a concentration of 0.1 g/mL buffer A supplemented with 1−1.5% (w/v) n-decyl-β-maltoside (DM, Glycon). Solubilized SLC26Dg was purified by immobilized metal affinity chromatography (IMAC). Target protein was immobilized on Ni-NTA resin and impurities were removed with 20 column volumes (CV) washing with 20 mM HEPES, pH 7.5, 150 mM NaCl (buffer B) supplemented with 50 mM imidazole, pH 7.5 and 0.2% DM. Protein was eluted with buffer B containing 300 mM imidazole and cleaved by HRV 3C protease during dialysis against buffer B without imidazole. Histidine-tagged GFP and protease were removed by IMAC, and cleaved protein was concentrated and subjected to size-exclusion chromatography (SEC) on a Superdex 200 Increase 10/300 column (GE Healthcare) equilibrated with 20 mM HEPES, pH 7.5, 150 mM NaCl and 0.2% DM (buffer C).

### Site directed spin labeling of SLC26Dg cysteine mutants

Cultivation and isolation of membrane vesicles were essentially performed as detailed above, but buffers for resuspending cells and membrane vesicles were supplemented with 3 mM, and 1 mM dithiothreitol (DTT), respectively. IMAC purification was conducted in the same way as described in the previous section but 5 mM 2-mercaptoethanol was included in all purification buffer to preserve the reduced state of the cysteine residues. Peak fractions from SEC purification were pooled and 2-mercaptoethanol was removed with Econo-Pac 10DG desalting column (Bio-rad) which was pre-equilibrated with buffer C. The concentration of eluted protein was adjusted to 7.5 μM with buffer C. The labeling of cysteine residue was initiated by stepwise addition of 100 mM MTSL spin label (in DMSO, Toronto Research Chemicals) in the protein solution to a final concentration of 300 μM and incubated at room temperature for 45 minutes with gentle agitation. The spin-labeled protein was further concentrated and free label was removed using Micro BioSpin 6 Chromatography Columns (Bio-rad) pre-equilibrated with buffer C.

### Reconstitution of SLC26Dg, SLC26Dg-IL and spin-labeled SLC26Dg variants

Proteoliposomes were essentially prepared as described previously^4, 49^. SEC-purified SLC26Dg and spin-labeled SLC26Dg variants in 0.2% DM was added to the liposomes (L-α-phosphatidylcholine from soybean) at a weight ratio of 1:50 (protein/lipid) for transport assays or 1:20 (protein/lipid) for PELDOR measurements, and detergent was subsequently removed by the addition of Biobeads. For radioisotope transport assays, proteoliposomes were harvested by centrifugation for 1.5 h at 250,000 × g and resuspended in 50 mM sodium phosphate (NaPi), pH 7.5, 2 mM MgSO4 to a lipid concentration of 20 mg/ml. After three freeze-thaw cycles, proteoliposomes were stored in liquid nitrogen until analysis. For PELDOR measurements, proteoliposomes were harvested by centrifugation for 20 min at 250,000 × g and resuspended in 50 mM KPi, pH 8.0 to a final spin concentration of 80 to 130 μM.

### PELDOR EPR

All the PELDOR experiments were performed at Q-band frequencies (33.7 GHz) using a Bruker E580 spectrometer equipped with an EN 5170 D2 cavity, 150 W traveling-wave tube (Applied Systems Engineering Inc.) microwave amplifier, and an ELEXSYS SuperQ-FT accessory unit. The temperature was kept at 50 K using a ITC 502 temperature control unit (Oxford Instruments) and a continuous-flow helium cryostat (CF935, Oxford Instruments). For all samples, 20 % (v/v) deuterated glycerol was added. For measurement, a 10 μL sample was transferred into a 1.6 mm outer diameter quartz EPR tubes (Suprasil, Wilmad LabGlass) and immediately frozen in liquid nitrogen. The dead-time free four-pulse PELDOR sequence with a phase-cycled π/2-pulse was used^50, 51^. Typical pulse lengths were 22 ns (π/2 and π) for the observer pulses and 12 ns (π) for the pump pulse. The delay between the first and second observer pulse was increased by 16 ns for eight steps to average deuterium modulations. The frequency of the pump pulse was set to the maximum of the echo-detected field swept spectrum to obtain maximum inversion efficiency. The observer frequency was set 70 MHz lower.

### CW EPR

Continuous wave (CW)-EPR spectra were recorded to determine the spin labeling efficiency. The spectra were recorded at a Bruker ELEXSYS E500 spectrometer (9.4 GHz) at room temperature with the following parameter settings: microwave power of 2.00 mW, modulation amplitude of 0.15 mT, and modulation frequency of 100 KHz.

### MD simulation of the SLC26Dg monomer

The crystal structure of the membrane domain of SLC26Dg monomer (PDB: 5DA0)^4^ was used in all-atom explicit solvent molecular dynamics (MD) simulation for equilibration and to uncover structural flexibility. The WT MD simulation model included residues Q14 to S392. The unresolved region between TM12 and TM13 (T334, L335, T336, V337) was modeled using Modeller^52^. The transmembrane domain of SLC26Dg was embedded into 241 palmitoyl-oleoyl-phosphatidylcholine (POPC) lipids and 13185 TIP3P water molecules^53^ were added (total system size 77650 atoms). We used GROMACS 5.1.3^54^ to perform simulations with a time step of 2 fs at a constant temperature (303.15 K) set with a Nosé-Hoover thermostat^55^ using a coupling constant of 1.0 ps. A semi-isotropic Parrinello-Rahman barostat^56^ was used to maintain a pressure of 1 bar. The all-atom CHARMM36 force-field was used for the simulation of protein and lipids^57, 58^. We performed the monomer MD simulation for around 1.1 μs.

### Structural modeling of the SLC26Dg dimer using PELDOR time traces

The conformation of monomeric SLC26Dg at 440 ns of the MD simulation was used for investigation of the dimeric state. We performed rigid body docking by placing a second protomer against the first protomer. We imposed C2-symmetry by rotating the second protomer in steps of 10 and 5 degrees about axes normal to the membrane plane centered at protomer two and one, respectively, and then bringing the two protomers to contact. For conformations without steric clashes, we forward calculated the PELDOR signals assuming uniform spin-label rotamer distributions^59^. We found that the interface had to be formed by TM13 and TM14 to match the PELDOR data for L385R1, which place the two sites in close spatial proximity.

### Structural modeling of the SLC26Dg dimer using PELDOR distance distributions

The SLC26Dg conformation at 440 ns of the MD simulation was used for rigid body docking using distance distributions (P(r)s as mean ± SD) and C2 symmetry as the restraints. Docking was performed using a grid search approach as implemented in the MMMDock tool of the Matlab-based software MMM^59, 60^. For more details see Supporting Information.

### MD simulation of the SLC26Dg dimer

To relax the SLC26Dg dimer conformation (docked based on PELDOR time traces), we performed an additional MD simulation. The dimer was embedded into 648 POPC lipids and 37227 TIP3P water molecules (total system size 210115 atoms). We used the same settings for the dimer MD simulation as described above for the monomer MD simulation. We performed the MD simulation for around 200 ns.

### BioEn spin-label reweighting

We calculated PELDOR signals for SLC26Dg dimer conformations saved along the MD simulation at 1-ns intervals. For each saved conformation, we performed a spin-label rotamer refinement^35^ by (1) attaching MTSL labels^59^, (2) calculating PELDOR traces for each label position (K353, V367, L385) and rotamer combination, and (3) ensemble-reweighting the spin-label rotamers using the “Bayesian inference of ensembles” (BioEn)^61, 62^ maximum-entropy method for each individual dimer conformation. We selected the conformation at 107 ns as the SLC26Dg dimer conformation with minimal total *χ^2^* for further analysis. L-curve analysis was used to identify suitable confidence parameters *θ_i_* that trade off consistency between the simulated data at each site and experiment (using a chi-squared metric *χ^2^)* and the changes in the ensemble weights (using relative entropy 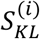 for label positions *i* = 1,2,3)^35^. Corresponding marginalized reweighted rotamer weights are visualized in Suppl. fig 4.

### Radioisotope transport assays

For transport studies, proteoliposomes were thawed and extruded through a 400 nm polycarbonate filter. Extruded proteoliposomes were pelleted by centrifugation for 20 min at 250,000 × g at 15 °C and resuspended to a final lipid concentration of 100 mg/ml in 50 mM NaPi, pH 7.5, 2 mM MgSO4. The sample was homogenized with a 26 gauge needle and stored at room temperature until use. Radioisotope transport studies were performed on stirred samples at 30 °C. To initiate transport, proteoliposomes were diluted 40-fold into the external buffer (50 mM KPi, pH 6.0, 2 mM MgSO4 containing 24 μM of [^14^C]-fumarate (Moravek) and 100 nM valinomycin). At appropriate time points, 100 μL samples were taken and immediately diluted with 2 mL ice-cold external buffer, followed by rapid filtration on 0.45 μm nitrocellulose filters. After washing the filters with another 2 mL buffer, the radioactivity associated with the filter was determined by scintillation counting.

### Oxidative crosslinking of SLC26Dg-IL

SLC26Dg-IL (I45C/A142C, inward-locked) was expressed and purified as described method for cysteine variants used in PELDOR studies. The eluted and HRV 3C protease-cleaved protein was subjected to SEC in buffer C supplemented with 3 mM DTT. Proteins from peak fractions were pooled and DTT was removed with an Econo-Pac 10DG desalting column (Bio-rad) which was pre-equilibrated with buffer C. The concentration of protein was adjusted to ~7 μM and oxidative crosslinking was initiated by adding a 10-fold concentrated CuPhen stock (3 mM CuSO4, 9 mM 1,10-phenanthroline monohydrate, freshly prepared) into the protein solution. The sample was incubated at room temperature for 45 min with gentle agitation and subsequently 0.5 M Na-EDTA, pH 7.0 was added to a final concentration of 20 mM to quench the reaction. The locked protein was injected for SEC to remove the crosslinking reagent and peak fractions were pooled and used for subsequent reconstitution and transport studies.

### Oxidative crosslinking of cysteine scanning mutants along TM14

*E. coli* MC1061 containing pBXC3sfGH holding the gene coding for cysteine variants of SLC26Dg fused C-terminally to superfolder GFP^63^ were cultivated in 700 μL of TB/Amp in a 96 deep well plate. Cells were grown at 37 °C until an OD_600_ ≈ 1 was reached, after which the temperature was gradually decreased to 25 °C over the course of 1 h. Expression was induced by the addition of 0. 005% L-arabinose and proceeded for 16 h. Cells were pelleted and resuspended in 500 μL of 50 mM KPi, pH 7.5, 1 mM MgSO4, 10% glycerol, 3 mM DTT, 1 mM PMSF with trace amounts of DNase I. After 20 min incubation on ice, cell disruption was carried out by 1 min sonification on ice at output level 4, and a 50% duty cycle (Sonifier 250, Branson). Unbroken cells and debris were remove by 5 min centrifugation at 13,000 × g. The supernatant was collected and DTT was removed by a Bio-Spin 6 column (Bio-rad) which was pre-equilibrated with 50 mM NaPi, pH 7.2. Oxidative crosslinking was initiated by adding a 10-fold concentrated CuPhen stock (3 mM CuSO4, 9 mM 1,10-Phenanthroline monohydrate, freshly prepared) into the vesicle solution. Samples were incubated at room temperature for 20 min and terminated by adding 100 mM NEM and 0.5 M Na-EDTA, pH 7.0 in a final concentration of 5 mM and 20 mM, respectively. The reaction mixture was mixed with SDS-PAGE sample buffer and the degree of crosslinking was determined by 8% SDS-PAGE and in gel GFP fluorescence imaging (ImageQuant LAS4000).

### Preparation of CHO cell membrane vesicles

CHO cells were cultivated as described previously^64^. 24 to 36 h after transfection, CHO cells expressing eGFP-tagged rat prestin (cysteine free) or single-cysteine variants (V499C and I500C located at the cytoplasmic end of TM14^65^) were treated with trypsin, harvested and washed with PBS containing 10 mM DTT. The cells were resuspended in disruption buffer (20 mM HEPES, pH 7.5, 150 mM NaCl, 10% glycerol, 1 mM MgSO4, 3 mM DTT, 20 μg/mL DNase I, 1 tablet of EDTA-free protease inhibitor cocktail) and lysed by 6 series of 5 sec sonification at 100% amplitude with 1 min incubation on ice in between (Sonoplus GM mini 20, Bandelin). The unbroken cells were removed by centrifugation at 1500 × g for 5 minutes and the membranes were obtained by ultracentrifugation at 50,000 rpm for 1 hour (TLA110 rotor). The membrane pellet was resuspended in membrane resuspension buffer (20 mM HEPES, pH 7.5, 150 mM NaCl, 10% glycerol, 3 mM DTT) to a final protein concentration of ~0.7 mg/mL.

### Oxidative crosslinking of CHO cell membrane vesicles

The resuspended rat prestin membrane vesicles were pelleted at 200,000 × g for 30 min at 4°C. After centrifugation, the membrane vesicles were gently washed with 20 mM HEPES, pH 7.5, 150 mM NaCl, 10% glycerol without disturbing the pellet. The crosslinking reaction was initiated by homogeneously resuspension of membrane vesicles in the same amount of crosslinking buffer (20 mM HEPES, pH 7.5, 150 mM NaCl, 10% glycerol, 300 μM CuSO4, 900 μM 1,10-phenanthroline monohydrate, freshly prepared) and incubated at room temperature for 20 min. The crosslinking reaction was quenched by addition of 0.5 M Na-EDTA to a final concentration of 20 mM. To decrease anomalous migration of rat prestin in SDS-PAGE, crosslinked vesicles were treated with PNGaseF (New England BioLabs) at 37°C for 1.5 hr. The reaction mixture was mixed with non-reducing SDS-PAGE sample buffer and the degree of crosslinking was determined by 8% SDS-PAGE and in gel GFP fluorescence imaging (ImageQuant LAS4000)

## Acknowledgements

We acknowledge Prof. Thomas Prisner for his support on the EPR measurements. We thank Melanie Engelin and Kai Steinmetz for their support in mutagenesis. The authors acknowledge financial support from the Max Planck Society (GH), the International Max Planck Research School (IMPRS) of the MPI of Biophysics in Frankfurt (EAJ), and the German Research Foundation via the Cluster of Excellence Frankfurt (Macromolecular Complexes; GH, ERG), the SPP1608 (Ultrafast and temporally precise information processing: normal and dysfunctional hearing; OL 240/4-2 to DO), and the SFB807 (Transport and Communication across Biological Membranes; GH, BJ, ERG).

## Supplementary methods

### PELDOR distance based rigid body docking

To determine the initial dimer structure model, rigid body docking was performed using the MMMDock tool in the Matlab based software MMM. The experimental distance distributions (P(r)s) for the positions K353R1 (4.4 ± 0.2 nm), V367R1 (3.9 ± 0.3 nm), and L385R1 (1.8 ± 0.1 nm) were used as the constraints, because the corresponding PELDOR time traces show clear oscillations. Since the STAS domain of the SLC26Dg monomer is in a non-physiological orientation in the crystal structure, the STAS domain was deleted and the MD refined monomer was used as the starting structure. Assuming C2 symmetry for the dimer (γ = 0) and a parallel orientation (z = 0), a grid of the angle values for α (between 0-360° with 10° steps) and β (between 0-180° with 5° steps) and of the translation parameters x and y (between ±7.5 nm with 0.25 nm steps) was generated.

For each model corresponding to a particular parameter set (α_i_, β_i_, x_i_, y_i_), mean distances for the restraints K353R1, V367R1, and L385R1 were simulated. To obtain the initial grid search dimer model, the model with the minimum root mean square deviation (RMSD) to the input values was chosen. The parameter set from this initial search (α_0_ = 360°, β_0_ = 5°, x_0_ = 5 nm, y_0_ = 1 nm) served as the starting parameters for subsequent refinement, where small changes of the parameters further minimized the RMSD. Since the refinement does not sample the whole possible parameter space, it was used after a global grid search to avoid being caught in local minima of the error surface. The parameters for the final model are α = 359.93°, β = 5.49°, x = 5.034 nm, y = 1.022 nm.

---

**Suppl. Fig. 1.**
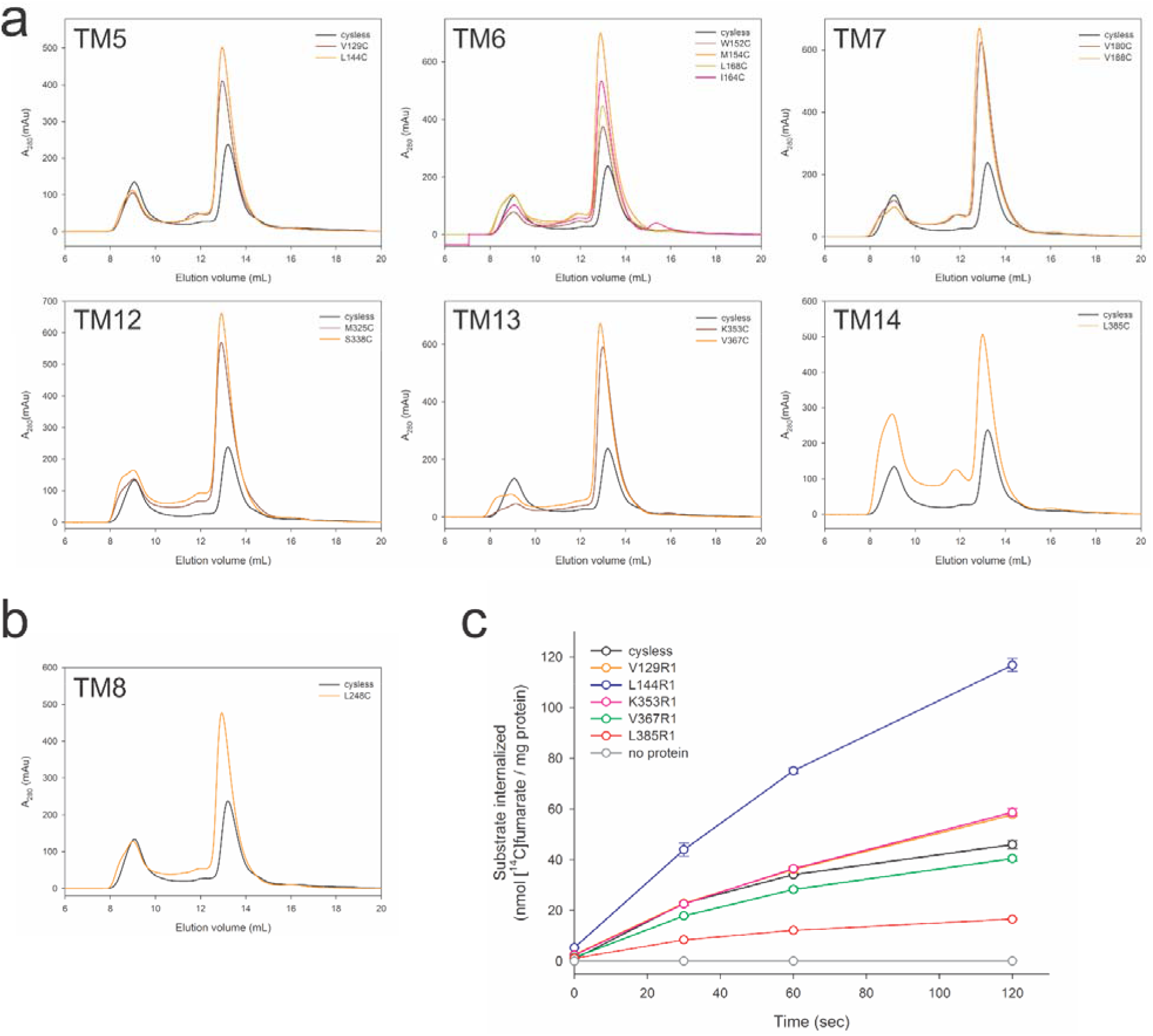
Assessment of the folding state of single cysteine mutants of SLC26Dg. **(a)** Analysis of decylmaltoside-solubilized gate domain cysteine mutants of SLC26Dg by size exclusion chromatography. **(b)** Analysis of decylmaltoside-solubilized core domain mutant L248C by size exclusion chromatography. **(c)** Fumarate transport of spin-labeled SLC26Dg mutants in proteoliposomes. Substrate transport was normalized to the amount of membrane-inserted protein. Data represent mean and standard errors of three technical replicates.

**Suppl. Fig. 2.**
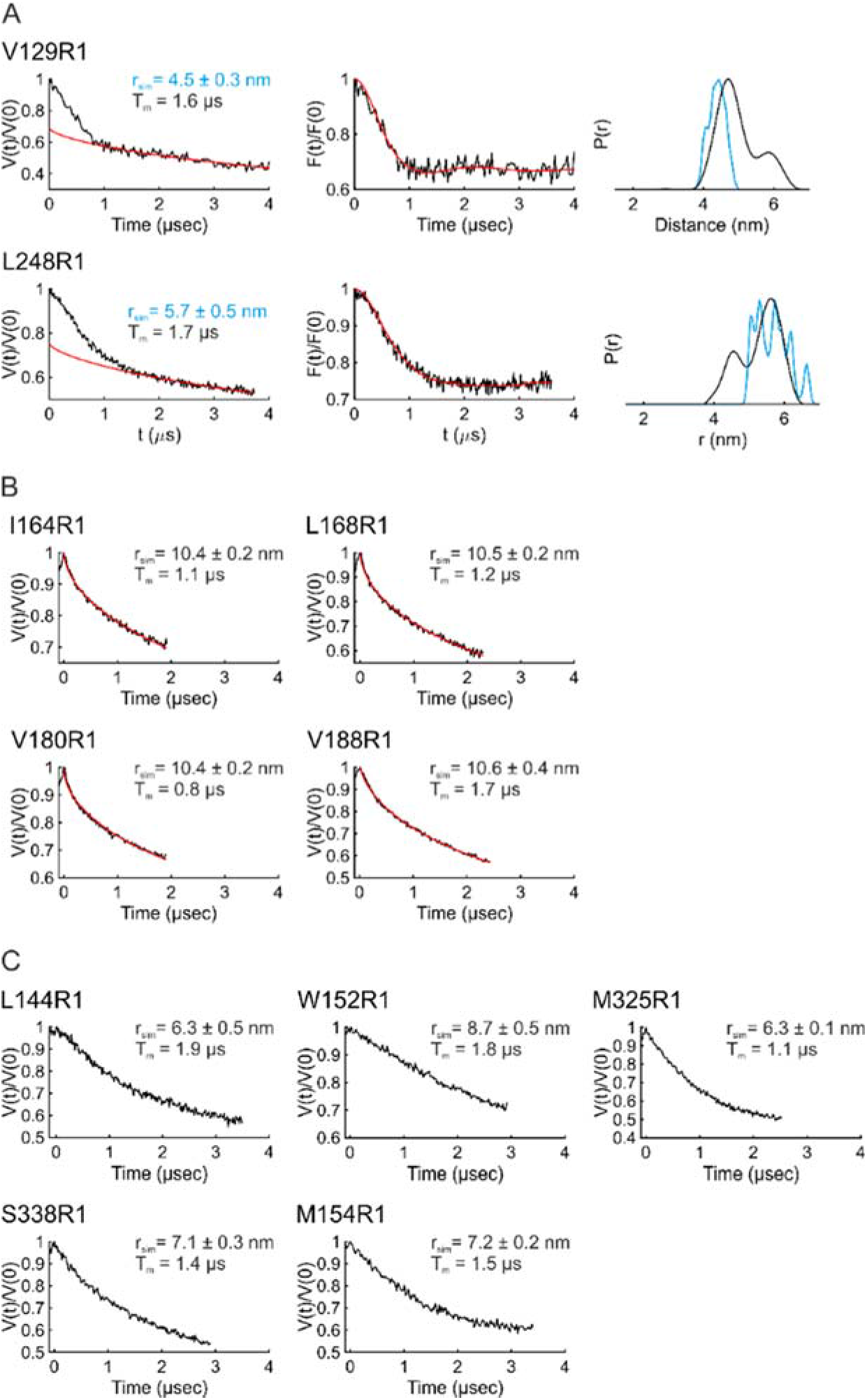
Overview of PELDOR measurements on membrane-reconstituted SLC26Dg. Experimental signal traces (black lines) are shown, where available, together with simulations (red) for the dimer model (using the form factors) and P(r) distance distributions (for V129R1 and L248R1). The phase memory time (*T_M_*) is listed. All the positions revealed a rather short *T_M_*, which permitted reliable distance determinations for only few positions. Overall, the data revealed a two-dimensional distribution of the spins (with *d* = ~2.0), which is expected for liposome-reconstituted membrane proteins. **A**) Original data (left), background corrected form factor (middle, in black) overlaid with the fit from Tikhonov regularization (in red), and the distance distribution (right) for positions V129R1 located on TM5 (in the gate domain) and L248R1 located on TM8 (in the core domain). The corresponding simulations (for P(r), in blue) performed on the dimer model (generated using the form factors) are also shown. Overall, the experimental and simulated distances show rather good agreement and indicate some additional flexibility in proteoliposomes as compared to the static dimer model. **B**) Original data for positions on TM6 and TM7 (in the gate domain), all of which revealed only a stretched exponential decay (d = ~2.0) without well-defined characteristic length. For these positions, simulations predict interspin distances >10 nm, which already might fall into the intermolecular distance regime and therefore cannot be separated anymore. **C**) For positions L144R1 (TM5), W152R1, M154R1 (TM6), M325R1 and S338R1 (TM12), simulations reveal distances between 6.3 and 8.7 nm. Although the PELDOR data reveal the presence of inter-protomer distances, owing to short *T_M_* the data could not be acquired long enough to determine those distances. In summary, positions V129R1 and L248R1 directly and positions I164R1, L168R1, V180R1, and V188R1 indirectly validate the SLC26Dg dimer model.

**Suppl. Fig. 3.**
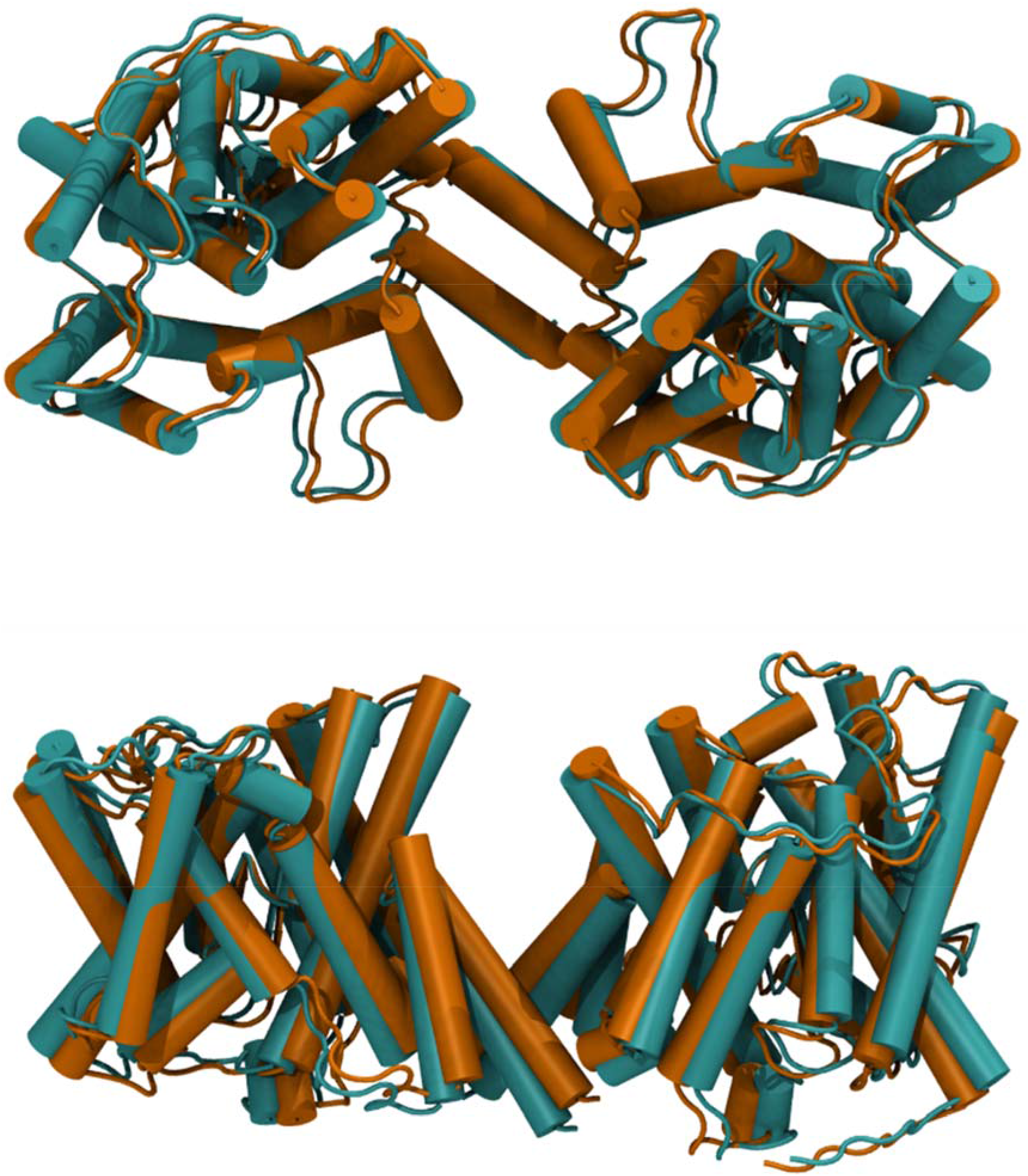
Comparison of SLC26Dg dimer models obtained via independent approaches. Superimposition of the models generated using MMMDock (orange) based on distance distributions P(r) and the model based on form factors calculated for MD dimer structures using BioEn rotamer refinement (teal). View from the extracellular side (top) and in the plane of the membrane (bottom). Neither model has backbone clashes. The BioEn model does not have any side chain clashes either. The two models align to each other with a global backbone RMSD of 1.7 Å. The MMMDock model exhibits two side chain clashes as well as contacts between some of the side chain atoms. The form factor based structure which is further minimized with MD simulation is the structure discussed in the manuscript. Despite the small differences between the structures, the fact that a very similar structure is obtained via two independent approaches largely excludes any bias by the fitting routines for the dimer structure.

**Suppl. Fig. 4.**
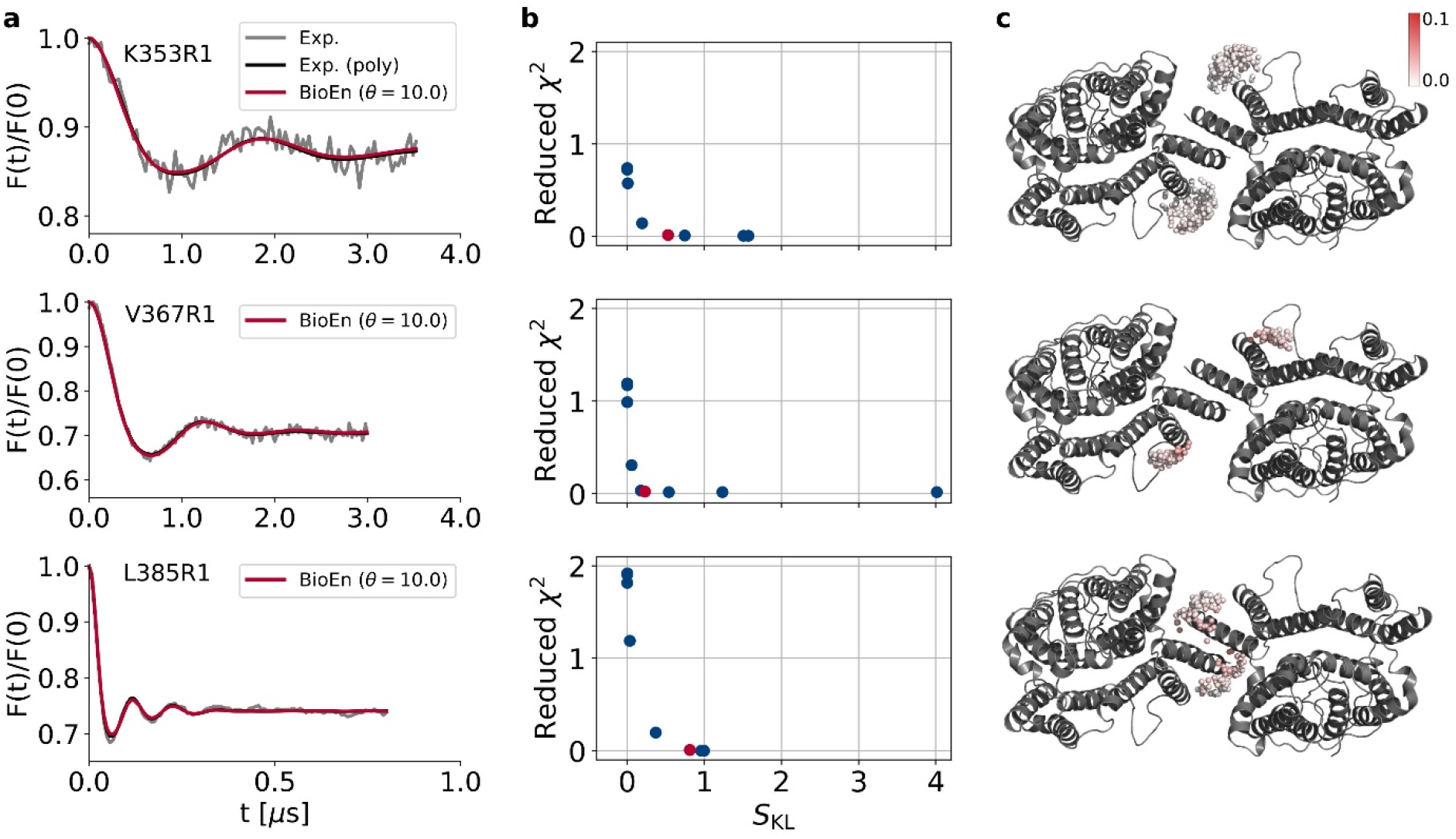
BioEn spin-label rotamer refinement using experimental PELDOR signals reveal weights of rotamer states. **(a)** For each labeled position (K353R1, V367R1, L385R1), the experimental PELDOR signals (gray), a polynomial fit to the experimental signals (black) and the calculated BioEn PELDOR traces after spin-label ensemble refinement are shown. Within the BioEn procedure, for each calculated PELDOR signal the result of the optimal confidence value *θ_i_* is shown. **(b)** L-curve analysis for the three PELDOR measurements. Reduced *χ^2^* is plotted as a function of the Kullback-Leibler relative entropy *S_KL_* for *θ =* 10^6^ to *θ* = 10^−1^ (left to right; blue), with *θ* decreased stepwise by factors of 10. The rightmost point is the result of a regular *χ^2^* minimization *(θ =* 0). Red circles indicate the results for *θ =* 10^1^, which is the value of *θ* chosen for further analyses. (c) Spin-label centers for SLC26Dg dimer (white to red spheres). Darker colors indicate higher relative weights (see scale bar).

**Suppl. Fig. 5.**
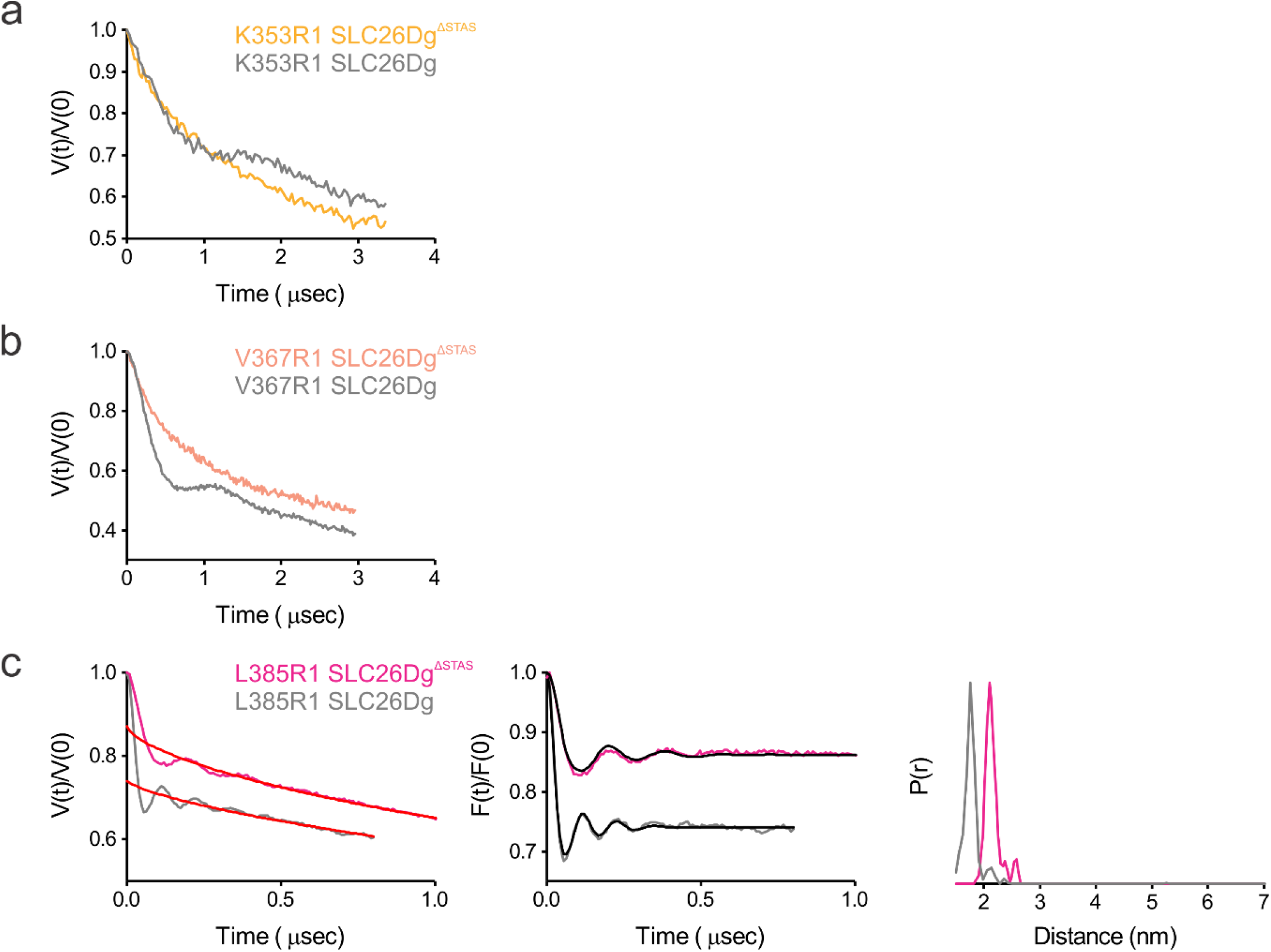
Interspin distances in the SLC26Dg^ΔSTAS^ dimer. Comparison of PELDOR data for membrane-reconstituted full-length (grey) and STAS deletion mutants of SLC26Dg as indicated. **(a-b)** For positions K353R1 and V367R1, which are located on TM13, deletion of the STAS domain results in an increased flexibility as reflected by the faster damping of V(t)/V(0) and by the presence of additional frequencies, corresponding to longer distances. Due to the insufficient time window for these SLC26Dg^ΔSTAS^ mutants, only a comparison of the original data is shown. **(c)** For L385R1 located on TM14, a small shift of the mean distance is observed. Altogether, the data show that STAS deletion does not prevent dimerization.

**Suppl. Fig. 6.**
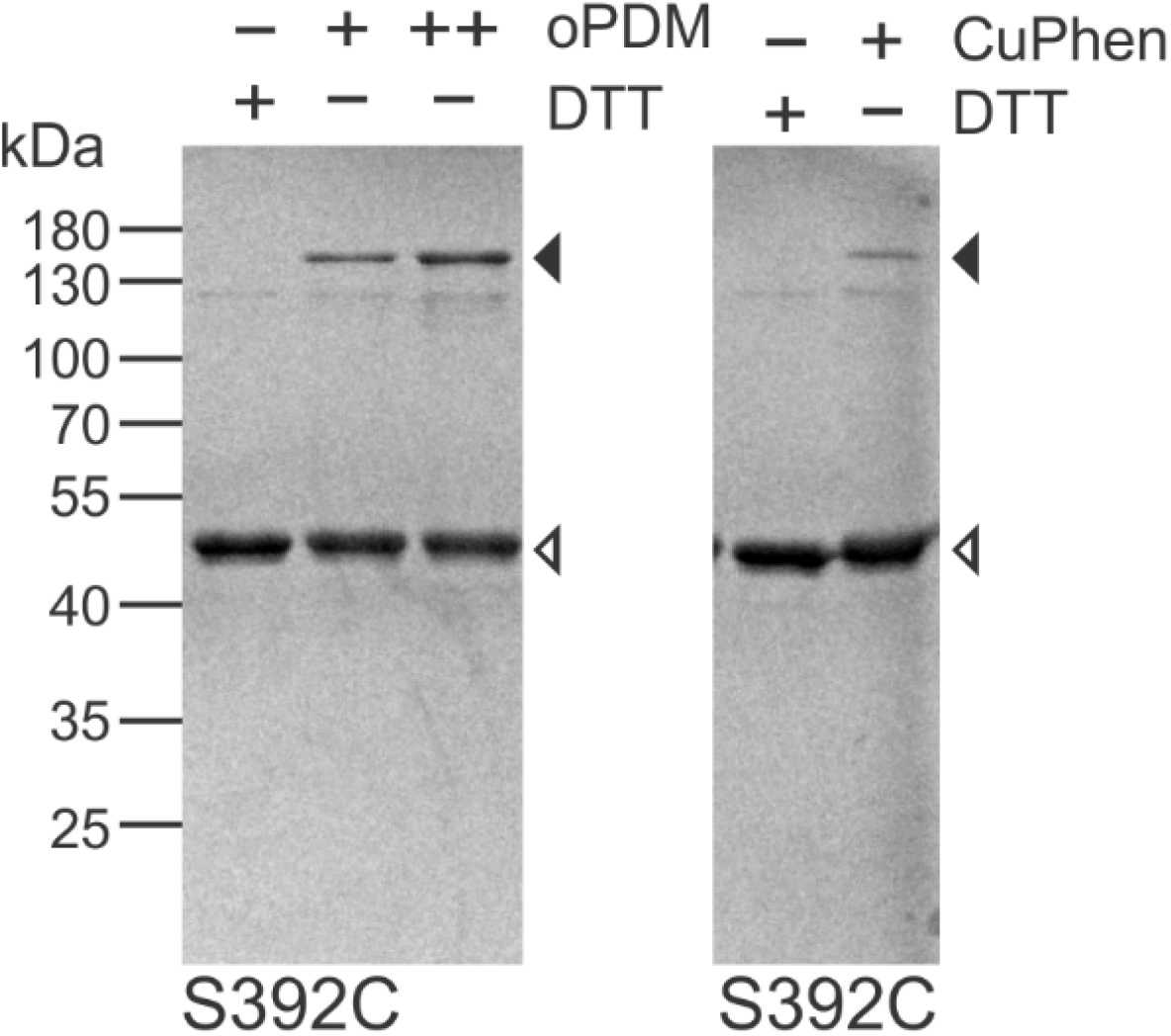
Cysteine crosslinking of SLC26Dg(S392C) in proteoliposomes. **(a)** SLC26Dg(S392C) was purified and membrane-reconstituted in the presence of reducing reagents. Crosslinking was initiated by the addition of 50 μM (“+”) or 500 μM (“++”) N,N’-1,2-phenylenedimaleimide (oPDM, left panel) or Cu-phenanthroline (right panel). Samples were incubated for 20 min at room temperature and subsequently analyzed by SDS-PAGE and visualized by coomassie staining. Black and white arrows indicate dimeric and monomeric SLC26Dg, respectively

**Suppl. Fig. 7.**
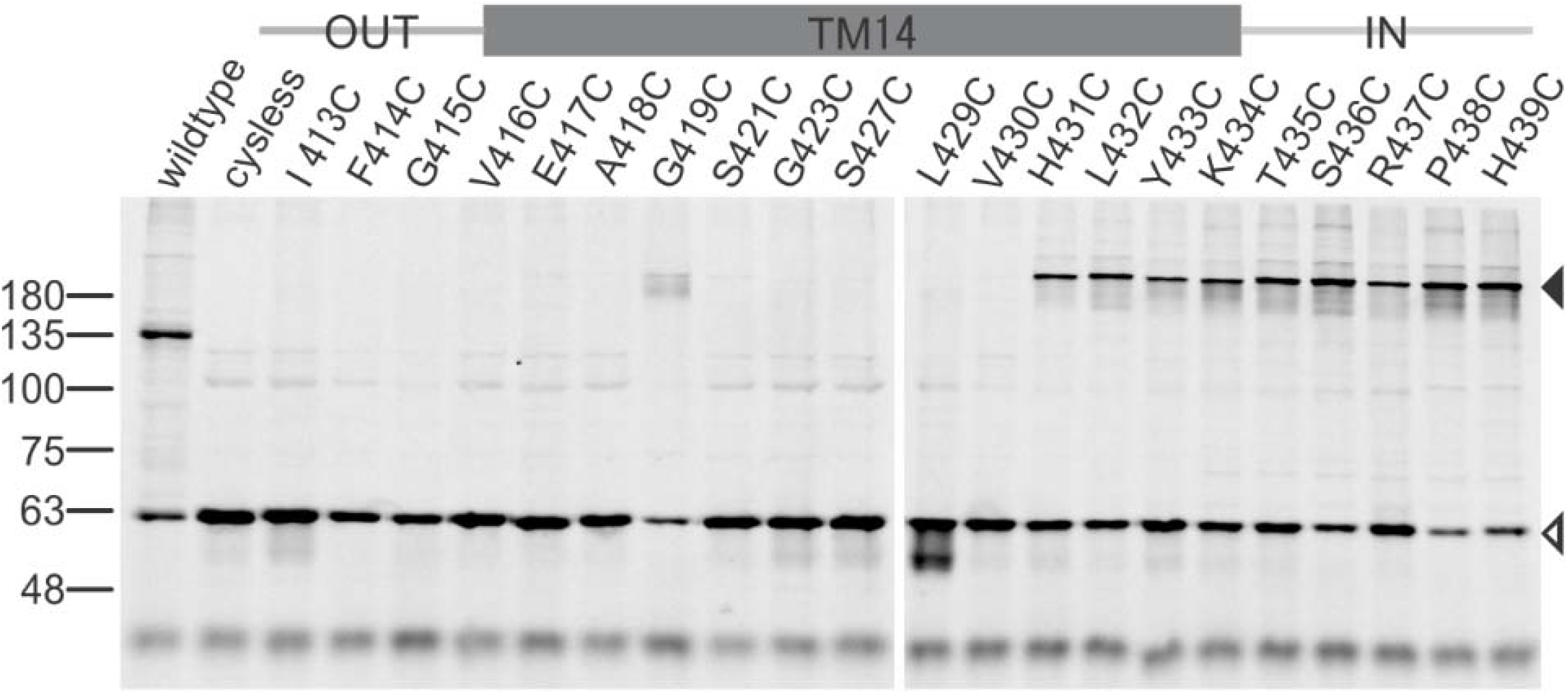
Oxidative cysteine crosslinking between TM14 of SLC26Si. In gel GFP fluorescence analysis of disrupted *E. coli* cells expressing single cysteine variants of SLC26Si fused to superfolder-GFP. The approximate boundaries of TM14 are indicated above. At the center of TM14, covering G419-L429, every second residue was analyzed. Following oxidative crosslinking, samples were analyzed by non-reducing SDS-PAGE. Wildtype (holding a native cysteine following the C-terminal STAS domain) and cysless represent positive and negative controls, respecti>vely. Black and white arrows indicate dimeric and monomeric SLC26Si, respectively.

**Suppl. Fig. 8.**
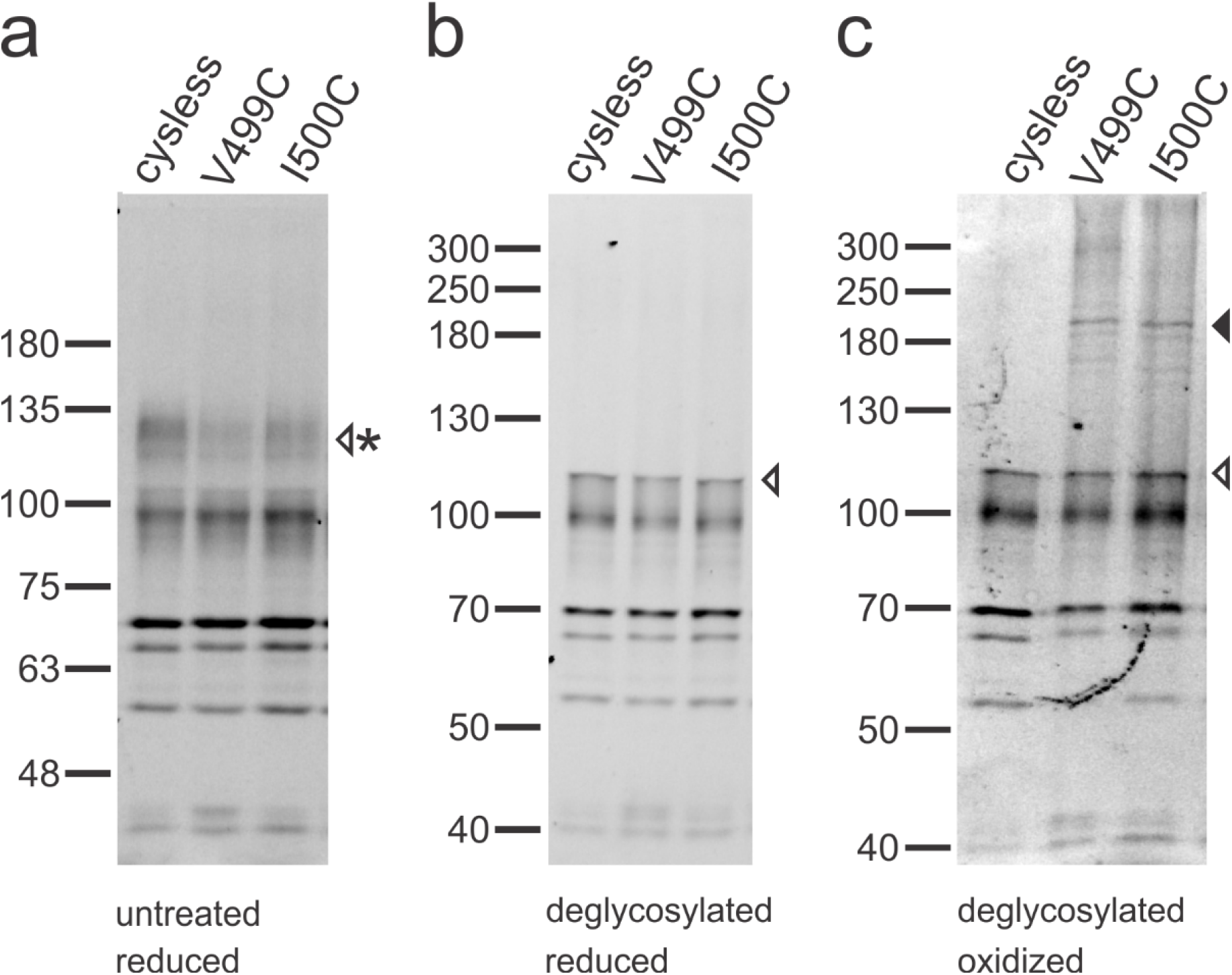
Oxidative crosslinking between TM14 of rat prestin. In gel GFP fluorescence analysis of CHO membrane vesicles holding cysteine variants of rat prestin fused to eGFP. V499 and I500 are located at the cytoplasmic end of TM14. Samples were analyzed **(a)** prior, or **(b)** following deglycosylation by reducing SDS-PAGE, or **(c)** following deglycosylation and oxidative crosslinking by non-reducing SDS-PAGE. Black and white arrows indicate dimeric and monomeric rat prestin, respectively. The asterisk indicates glycosylated rat prestin. Lower molecular weight bands represent mostly nonglycosylated, proteolytic fragments of rat prestin.

**Suppl. Fig. 9.**
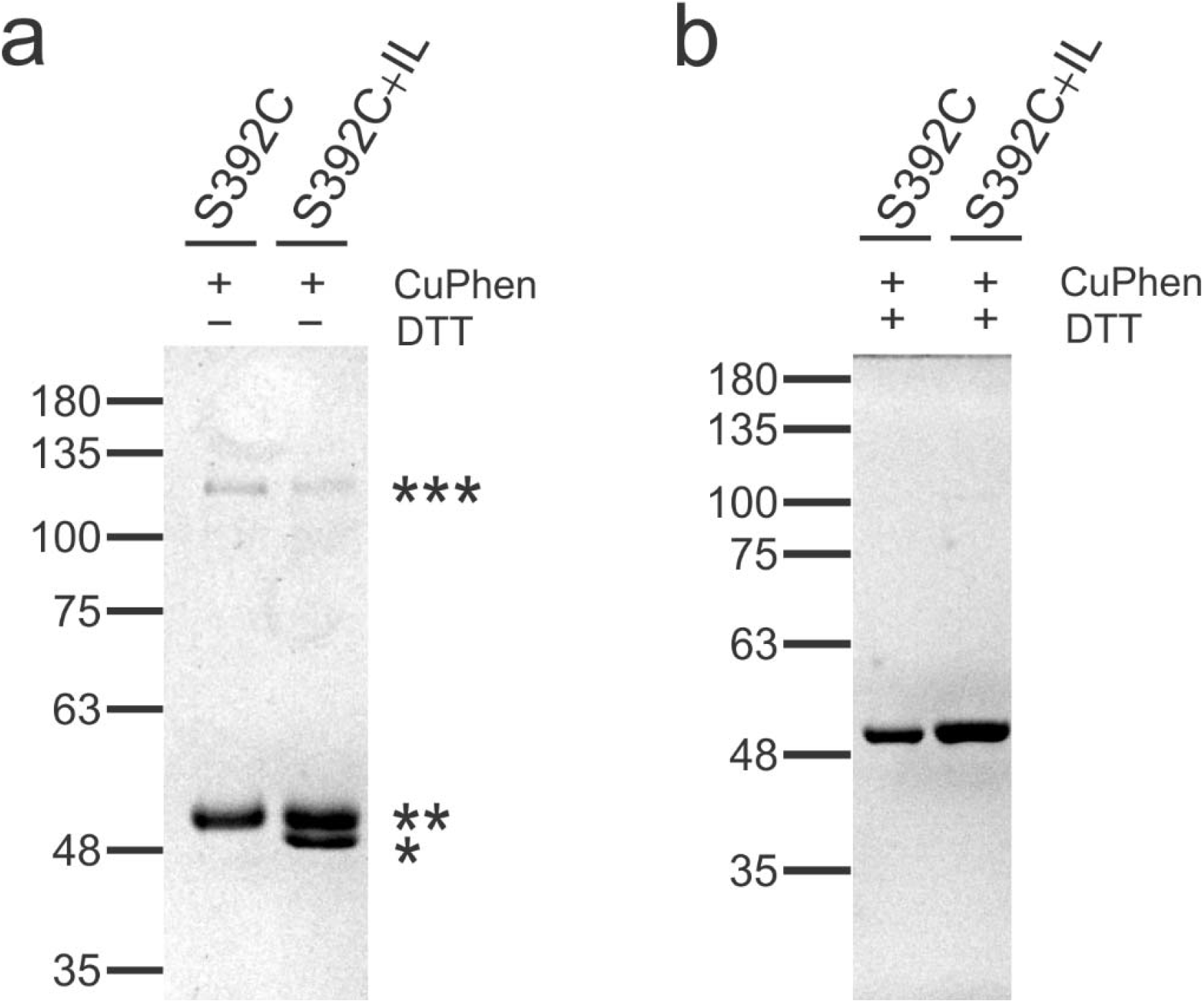
Stochastic formation of heterodimers upon membrane-reconstitution of monomers. **(a)** SLC26Dg(S392C) (100%) and a 50:50 mixture of SLC26Dg(S392C) : SLC26Dg-IL were membrane-reconstituted and following oxidative crosslinking, analyzed by non-reducing SDS-PAGE and visualized by coomassie staining. Similar amounts of S392C are loaded in both lanes. The triple, double and single asterisk indicate the S392C-S392C homodimer, the monomeric S392C and the crosslinked IL mutant, respectively. The decrease in the amount of crosslinked S392C upon addition of IL indicates the formation of S392C-IL heterodimers and suggests that protomers combine to dimers in a random manner. **(b)** Reducing SDS-PAGE analysis of samples from panel (a) visualized by coomassie staining.

**Suppl. Fig. 10.**
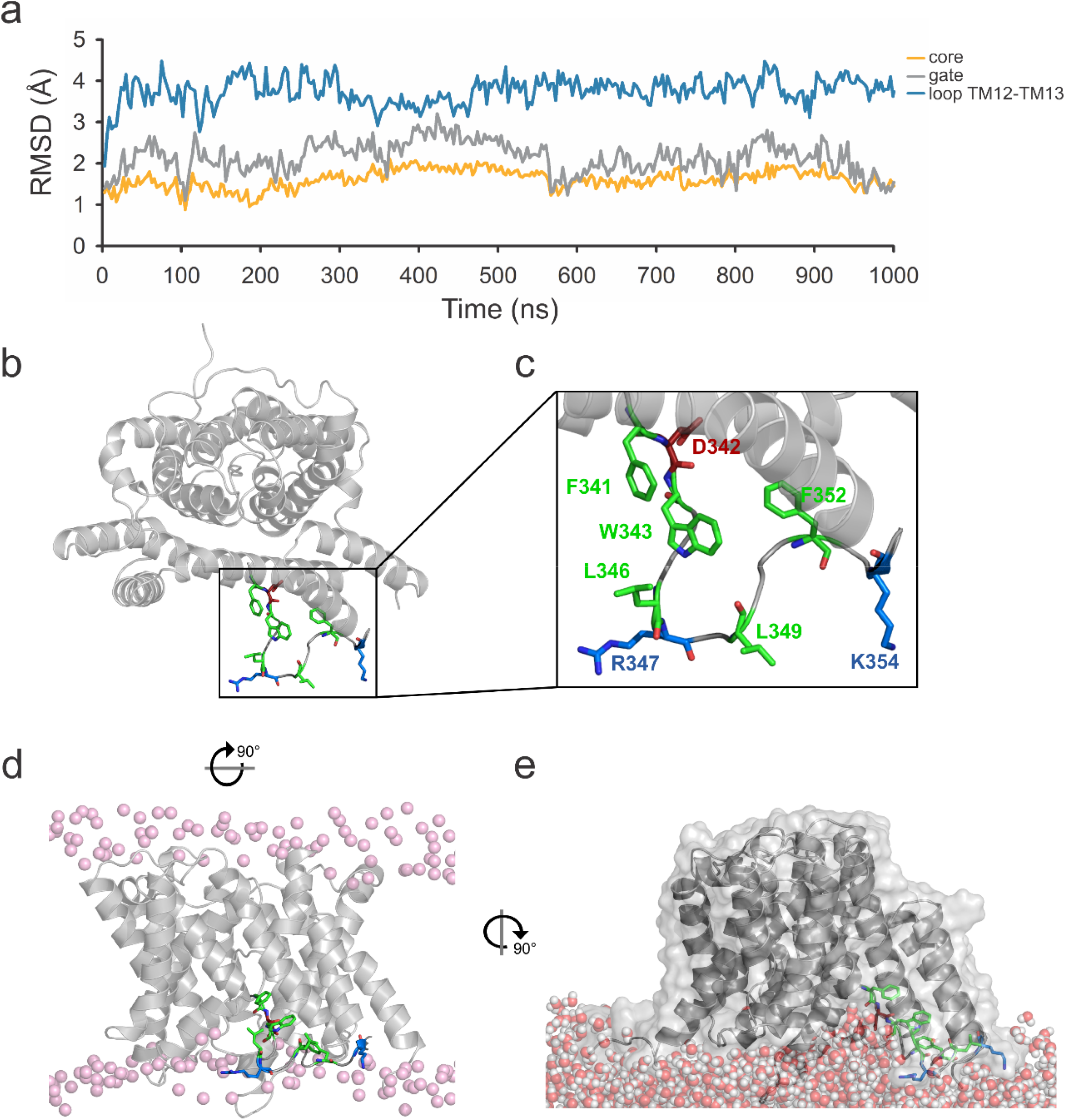
Reduced flexibility of the cytoplasmic loop connecting TM12 and 13 observed in the MD simulation. **(a)** C_α_-atom root-mean-squared-distance (RMSD) values of the core, gate, and the cytoplasmic loop connecting TM12 and 13 relative to the monomer crystal structure as a function of MD time (1 μs). **(b)** Top-view of SLC26Dg is shown with highlighted residues: R347 and K354 (blue), D342 (red) and hydrophobic residues F341, W343, L346, L349, F352 (green). **(c)** Detailed view on the residues of the respective. **(d)** Side-view of SLC26DG with residues highlighted and lipid headgroups (phosphorus atoms) shown as spheres. **(e)**. Turned side-view to indicate funnel of water molecules towards the binding site.

